# The Crimean population of the lesser grey shrike (*Lanius minor*) has low behavioural flexibility in its response to approaching humans

**DOI:** 10.1101/2022.07.02.498546

**Authors:** Peter Mikula, Zbigniew Kwieciński, Ireneusz Kaługa, Piotr Tryjanowski

## Abstract

The ongoing growth of the human population will increase the rate of wildlife−human interactions. High levels of animal tolerance and flexible responses towards human presence seem to be among the key mechanisms behind successful wildlife−human coexistence, but this behaviour remains unexplored for most populations and species of animals. Here, we investigate the escape behaviour (measured as flight initiation distance) of the Crimean population of a charismatic and declining bird species, the lesser grey shrike (*Lanius minor*). We examined its relationship with starting distance of the approaching human, directness of that approach (direct or tangential), habitat type (rural or suburban), and height of the perch used by shrikes. We found that the starting distance was significantly associated with escape responses of shrikes to approaching humans. In contrast, we found no significant association between escape responses and directness of approach, habitat type, or height of perch. Our results indicate that the lesser grey shrike may exhibit low flexibility in their escape responses towards humans which may have implications for their conservation management. Our results also indicate that the widely used 30 metres threshold for minimum starting distance may be insufficient for rural populations, even of small passerines.

## Introduction

The human population and its impact on natural ecosystems are both predicted to further increase during the 21^st^ century (Sanderson *et al*. 2002). Animals may adaptively respond to changed environmental conditions by cognitive processes adjusting their behaviour (Lima & Dill 1990; Ducatez *et al*. 2020). Although most wildlife−human interactions are non-lethal and harmless, animals usually perceive humans as predators and respond to them in the same way, including escaping (Frid & Dill 2002). Increased human presence may increase the frequency of costly escape responses in animals that lack the ability to tolerate human disturbance, with possible cascading effects on individual fitness and population dynamics (Bélanger & Bedard 1990; Steven *et al*. 2011; Møller *et al*. 2014; Moss *et al*. 2014). For example, European and Australian bird species with declining populations are less tolerant of an approaching human than species with increasing populations (Møller *et al*. 2014). Escape behaviour of animals is affected by several factors, such as the distance at which an approach begins (Blumstein 2003; Tryjanowski *et al*. 2020; Mikula *et al*. 2021), body mass and overall pace-of-life strategy (Stankowich & Blumstein 2005; Weston *et al*. 2012), sex and age (Kalb *et al*. 2019), flock size (Morelli *et al*. 2019; Tryjanowski *et al*. 2020), but also more subtle risk-cues such as human gaze and head orientation (Bateman & Fleming 2011; Clucas *et al*. 2013), and directness (Møller & Tryjanowski 2014) and speed of approach (Stankowich & Blumstein 2005). Based on their traits, some species are expected to have decreased ability to cope with human disturbance and the identification of these may help in their conservation management. However, the flexibility in behavioural responses towards human presence remains unexplored for many populations and species of animals.

The lesser grey shrike (*Lanius minor*) is a charismatic and emblematic species of passerine bird (Order: Passeriformes) (Fig. 1a) breeding in large areas from Spain to Central Russia. However, the population of this species has undergone a sharp decline in most European countries since the 19^th^ century; the species is currently considered to be extinct in several Western (e.g., Belgium and Germany) and Central (e.g., Czech Republic and Poland) European countries with only isolated populations surviving in others (e.g., Spain and Slovakia) (Bronskov & Keller 2020; Yosef & International Shrike Working Group 2020). This decline has been largely attributed to the abundance declines of its main diet type, i.e., large insects, mainly due to a general intensification of agriculture, including intensive use of insecticides (Bronskov & Keller 2020; BirdLife International 2022). It has been found that diet generalist species have higher technical innovation rates and larger brains, which may indicate lower behavioural flexibility and plasticity in diet specialist species such as the lesser grey shrike (Ducatez *et al*. 2015). However, there is virtually nothing known about the behaviour of these shrikes in the presence of humans (Díaz *et al*. 2013; Livezey *et al*. 2016). Here, we explore the flexibility of escape responses towards approaching humans of lesser grey shrikes inhabiting the Crimean Peninsula. We investigate the influence of starting distance of approaching human, type of approach (direct or tangential), habitat type (rural or suburban), and the height of the perch used by birds.

**Figure 1.**
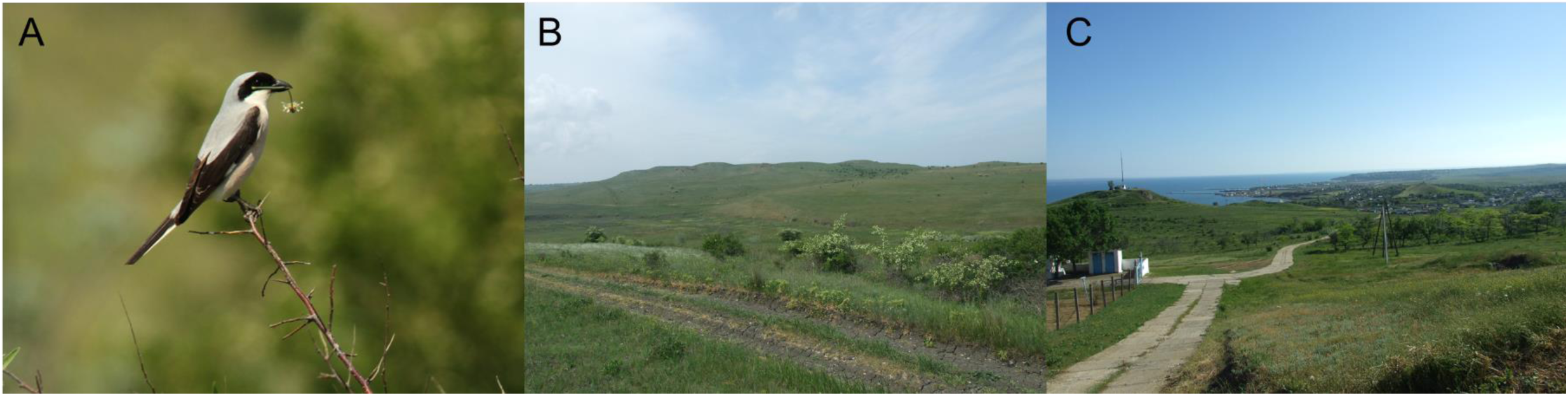
Photographs documenting (a) studied species, the lesser grey shrike (*Lanius minor*), and habitat types differing in the level of anthropogenic disturbance inhabited by this species in the Crimean Peninsula, including (b) rural (Beherove; 45°26′01″N, 36°14′55″E; 15 May 2013), and (c) suburban areas (Kercz lighthouse; 45°23′08″N, 36°38′13″E; 19 May 2013).

## Material and Methods

### Study area and data collection

Escape responses of lesser grey shrikes were collected in the Crimean Peninsula (*de jure* Ukraine) during May 2013. The study was conducted in the steppe landscape of the Kerch Peninsula, 20 km west of the city of Kerch (45°35′N, 36°46′E), covering six sampling sites: Zatyshe (45°22′28″N, 36°10′01″E), Chystopillya (45°22′01″N, 36°10′58″E), Beherove (45°26′01″N, 36°14′55″E), Oktyabrs′ke (45°21′48″N, 36°21′52″E), Kerch lighthouse (45°23′08″N, 36°38′13″E), and Kerch suburbs (45°17′48″N 36°25′19″E). Many natural hilly parts of the steppe have been preserved in this area. The study location (150 km^2^) is an extensively used landscape, mainly by grazing cattle. Generally, it is a grassland dominated by feather grass *Stipa capillata, S. pennata* and *S. pulcherrima*, furrowed fescue *Festuca sulcata* and dittany *Dictamnus albus*. The mosaic of this area is formed by midfield woodlots of different ages, scattered trees and discontinuous linear habitats, mainly consisting of mixed rows of shrubs and single trees such as Crimean pine *Pinus pallasiana*, Russian olive *Elaeagnus angustifolia*, false acacia *Robinia pseudoacacia*, and common hawthorn *Crataegus monogyna*. There are also seasonal streams, salt pans and mid-field ponds. The Crimean population of these shrikes is still quite numerous and the population appears stable (Andriushchenko & Vorovka 2022).

The escape responses of shrikes towards approaching humans were estimated by the flight initiation distance, a simple and widely adopted metric used as a proxy for fearfulness and willingness of animals to take a risk (Stankowich & Blumstein 2005; Blumstein 2006; Díaz *et al*. 2013). It has been shown that risk assessment measured by flight initiation distances is highly consistent for individuals, populations and species tested under similar circumstances (Carrete & Tella 2010, 2013; Díaz *et al*. 2013; Guay *et al*. 2016). In brief, all data were collected by a single author (Z. K.) wearing outdoor clothing (without any bright colours) using a standard procedure previously described (Blumstein 2006; Díaz *et al*. 2013). When a focal bird was spotted, the researcher moved at a normal walking speed (∼1 ms^-1^) either (a) directly towards the bird (N = 190 observations) or (b) tangentially towards the bird (i.e., passing the individual at a distance of ∼5 m) (N = 48; for similar approach, see Møller & Tryjanowski 2014). During both approaches, the head and gaze of the researcher was oriented towards the focal bird. The flight initiation distance was then estimated as the distance between the researcher (measured as a number of ∼1 m long steps) and the focal bird when it first started to escape (usually by flying away). If the bird was positioned above the ground, e.g., perching on vegetation, the flight initiation distance was estimated as the Euclidean distance between the position of the researcher and the focal bird. We examined only single bird individuals showing no initial signs of distress behaviour. We avoided re-sampling of the same individual by moving to a different shrike territory after each sampling occasion. All data were collected during favourable weather conditions (i.e., no rain, and no or light wind). In total, we collected 238 flight initiation distance estimates for shrikes (mean ± SD = 11.75 ± 6.04 metres).

For each observation, we also collected data on the starting distance, habitat type, and the height of perch used by shrikes. The starting distance was estimated as the distance (in metres) to the focal bird when the researcher initiated the approach (mean ± SD = 14.49 ± 8.34 metres). Habitat type was recorded as either rural (N = 143 observations; areas with little or no human population and buildings; Fig. 1b) or suburban (N = 95; typically areas on the edges of human settlements with 30−50% of built-up cover and >10/ha human population density; Fig. 1c) following the scheme by Marzluff *et al*. (2001). Perch height was estimated as the vertical distance (in metres) between the bird and the ground (recorded as zero if the bird was sitting on the ground) (mean ± SD = 3.08 ± 1.29 metres). All these factors have been previously found to affect avian responses towards approaching humans; escape distances typically increased with increasing starting distance (Blumstein 2003; Mikula *et al*. 2018; Tryjanowski *et al*. 2020), and from suburban to rural habitats (Samia *et al*. 2015b), but previous studies provided mixed results for the effect of perch height (Blumstein *et al*. 2004; Bjørvik *et al*. 2015). Each observation was also dated (so can also be used as a proxy for different sampling localities). We did not include the sex of birds in our analyses, because it was usually not possible to reliably sex birds in the field.

### Statistical analyses

All analyses were conducted in R v. 4.1.2 (R Development Core Team 2021).We tested for association between the flight initiation distance (response variable) and the starting distance (in metres; log10-transformed), approach type (direct or tangential), habitat type (rural or suburban), and height of the perch (in metres) (predictors) using a generalized linear model. The model was run using a Gaussian error distribution and a logarithmic link function. We graphically inspected the goodness of fit and the distribution of the residuals, revealing no major violations of the model assumptions. We calculated adjusted deviance explained by the model using the Dsquared function in modEvA v. 3.5 package (Barbosa *et al*. 2013). We checked collinearity between predictors using the vif function in car v. 3.0-10 package (Fox *et al*. 2020), revealing a low collinearity between predictors (<2 in all cases).

Because observed patterns could also be affected by sampling date and locality, we re-ran the same model using collection date (transformed to a categorical variable) as a random factor in a generalized linear mixed model using the lmer function in lme4 v. 1.1-27.1 and lmerTest v. 3.1-3 packages (Kuznetsova *et al*. 2013; Bates *et al*. 2015). Both models revealed the qualitatively same result, hence, we report only results using the simpler generalized linear model.

### Ethical note

Our study was observational and non-invasive. All fieldwork was conducted in accordance with approved guidelines. Data were collected in public places where no special permit was required. Our data collection caused only brief and minimal disturbance to birds which typically did not differ from standard background disturbance caused by other site visitors.

## Results

We found that only the starting distance significantly predicted shrike escape responses towards approaching humans (Table 1; Fig. 2). Although initial data exploration suggested that shrikes escaped significantly earlier in rural than suburban areas (ANOVA: F_1,236_ = 390.2, p <0.001), this was influenced by significantly longer starting distances in rural than suburban areas (ANOVA: F_1,236_ = 394.7, p <0.001). Moreover, we found that shrike escape responses did not differ significantly between direct and tangential approaches. Finally, escape responses of shrikes were not significantly correlated with perch height.

**Table 1.** Association between flight initiation distance (response variable) and starting distance (in metres; log10-transformed), approach type (direct or tangential), habitat type (rural or suburban), and height of the perch (in metres) (predictors) in the lesser grey shrike (*Lanius minor*) sampled in the Crimean Peninsula (N = 238 observations). The model explained 93% of the variation present in the data. Boldface indicates statistically significant results.

| Variable | Estimate | Standard error | t-value | p-value |
| --- | --- | --- | --- | --- |
| Intercept | 0.284 | 0.078 | 3.624 | <0.001 |
| **Starting distance** | **1.848** | **0.061** | **30.503** | **<0.001** |
| Approach (tangential) | 0.015 | 0.018 | 0.806 | 0.421 |
| Habitat (suburban) | -0.013 | 0.035 | -0.377 | 0.707 |
| Perch height | 0.017 | 0.009 | 1.936 | 0.054 |

**Figure 2.**
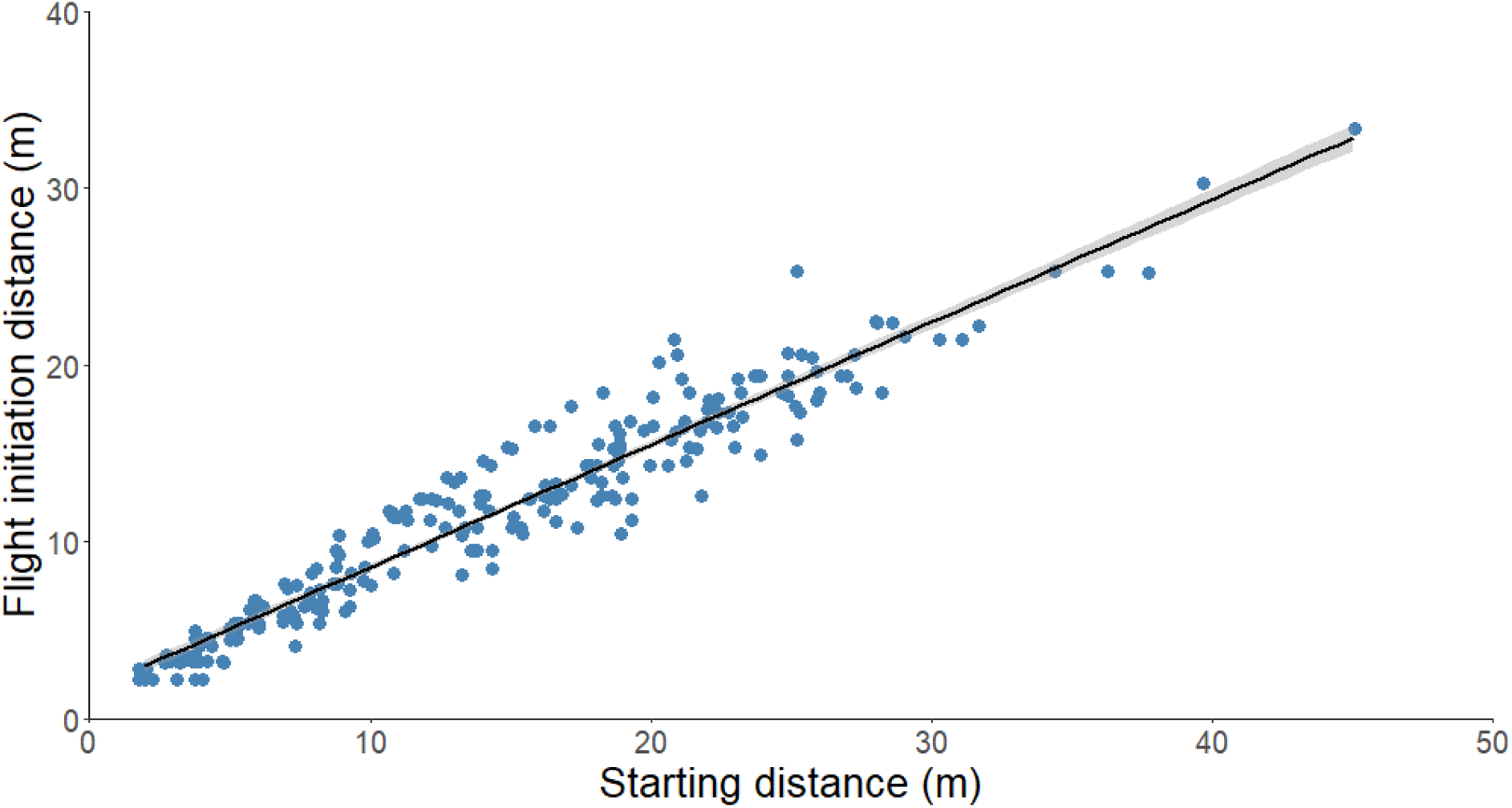
Association between flight initiation distance (response variable) and the starting distance (predictor) (original, untransformed values). Line and shaded area represent the results of univariate robust linear regressions and its 95% confidence interval (not controlled for other predictors) fitted using the rlm function in the MASS v. 7.3-55 package (Venables & Ripley 2002).

## Discussion

We found that escape distances in lesser grey shrikes occupying the Crimean Peninsula were significantly correlated only with starting distance. The directness of approach, habitat type, and perch height were not significantly correlated with their escape responses. This may indicate that the lesser grey shrike exhibited low flexibility in their response towards approaching humans which may have implications for the conservation of this species. This is interesting because shrikes are usually considered to be intelligent birds (Syrová *et al*. 2016) with a relatively large brain size (Sayol *et al*. 2016); larger-brained species were found to delay escape from predators (Samia *et al*. 2015a).

If not standardized, the starting distance, i.e., the distance from which an animal was first approached, is one of the strongest predictors of escape distances in birds (Blumstein 2003; Mikula *et al*. 2018; Tryjanowski *et al*. 2020). Birds may consider longer approaches as more dangerous or approaches from longer distances may require higher monitoring costs – to avoid such costs, birds may escape early after detection of an approaching threat (Blumstein 2010; Samia *et al*. 2013). A strong association between starting and escape distance has practical consequences for conservation management practices in this species. The lack of information on the starting distance may result in biased estimates of escape distances (Blumstein *et al*. 2015); various starting distances should be adopted to correctly estimate the maximum, the mean and the variance of escape distances in order to set up adequate buffer zones (Guay *et al*. 2016; Livezey *et al*. 2016). However, our results also indicate that estimating safe buffer zones from maximum escape distances may be a tricky task in the lesser grey shrike and other species that show no evidence of a plateau in the relationship between escape and starting distances, even if starting distances are long (> 40 metres in this small passerine) (Fig. 2). In these cases, rather than setting up excessively large buffer zones that may impede birdwatching and research activities, alternative strategies may be adopted such as forming permanent tourist paths and observation hides with larger buffers present mainly in sites with especially sensitive species. Finally, our results indicate that the widely used threshold for minimum starting distance that was argued to mitigate the effect of starting distance on flight initiation distance (30 metres; e.g., Díaz et al. 2013) may be insufficient even in small passerines outside urban areas.

Our findings suggest that the lesser grey shrike (at least, the studied Crimean population) exhibited relatively low behavioural flexibility in response to human approaches under various conditions. This may indicate that this species may be increasingly vulnerable to an increase in human activity. Further studies using additional populations of the lesser grey shrike, as well as other species, are required to address the question of whether habitat and diet specialist species are increasingly vulnerable to human disturbance. Finally, one of the main future challenges is the incorporation of these findings in efficient wildlife protection and management.

## Acknowledgements

We have no known conflict of interest to disclose. We would like to thank our Ukrainian and Russian hosts from The Azov–Black Sea Ornithological Station for help in the field. Special thanks to Dr. Yuriy Andriushchenko for agreeing to be our guide on the Kerch Peninsula. We would also like to thank Tim Sparks for English language correction.

## Author Contributions

P. Tryjanowski conceived the idea; Z. Kwieciński collected data in the field with assistance of I. Kaługa; P. Mikula analysed the data and wrote the manuscript with help from P. Tryjanowski and Z. Kwieciński. All authors approved the final version of the manuscript.

## Data Availability Statement

The data that support the findings of this study will be available in the FigShare Digital Repository after acceptance.

